# Rolling the Dice Twice: Evolving Reconstructed Ancient Proteins in Extant Organisms

**DOI:** 10.1101/027003

**Authors:** Betul Kacar

## Abstract

Scientists have access to artifacts of evolutionary history (namely, the fossil record and genomic sequences of living organisms) but they have limited means with which to infer the exact evolutionary events that occurred to produce today’s living world. An intriguing question to arise from this historical limitation is whether the evolutionary paths of organisms are dominated by internal or external controlled processes (i.e., Life as a factory) or whether they are inherently random and subject to completely different outcomes if repeated under identical conditions (i.e., Life as a casino parlor). Two experimental approaches, ancestral sequence reconstruction and experimental evolution with microorganisms, can be used to recapitulate ancient adaptive pathways and provide valuable insights into the mutational steps that constitute an organism’s genetic heritage. Ancestral sequence reconstruction follows a backwards-from-present-day strategy in which various ancestral forms of a modern gene or protein are reconstructed and then studied mechanistically. Experimental evolution, by contrast, follows a forward-from-present day strategy in which microbial populations are evolved in the laboratory under defined conditions in which their evolutionary paths may be closely monitored. Here I describe a novel hybrid of these two methods, in which synthetic components constructed from inferred ancestral gene or protein sequences are placed into the genomes of modern organisms that are then experimentally evolved. Through this system, we aim to establish the comparative study of ancient phenotypes as a novel, statistically rigorous methodology with which to explore the respective impacts of biophysics and chance in evolution within the scope of the Extended Synthesis.

---

Living organisms are historical systems and their evolutionary history is one of the vital determinants of their capacity to respond to their environment. Historical contingency is a property of living systems that assigns past events as important factors that shape a system’s current state. In his book *Wonderful Life,* Stephen Jay Gould famously posed a thought experiment to address whether current biota are the product of random evolutionary luck, or whether there were other possible trajectories that could have taken place. Assigning historical contingency as a fundamental influence in shaping evolutionary outcomes, Gould posited that if life’s tape were rewound and replayed from various points in the distant past, the resulting living world would be very different than it is now:

> I call this experiment “replaying life’s tape.” You press the rewind button and, making sure you thoroughly erase everything that actually happens, go back to any time and place in the past—say, to the seas of the Burgess Shale. Then let the tape run again and see if the repetition looks at all like the original. If each replay strongly resembles life’s actual pathway, then we must conclude that what really happened pretty much had to occur. But suppose that the experimental versions all yield sensible results strikingly different from the actual history of life? What could we then say about the predictability of self-conscious intelligence? or of mammals? or of vertebrates? or of life on land? or simply of multicellular persistence for 600 million years? (Gould 1989)

Understanding to what degree historical contingency shapes evolutionary trajectories would be possible by performing Gould’s thought experiment. We cannot, of course, carry out this experiment at the global scale Gould envisioned, however, methods of experimental evolution allow researchers to tackle some aspects of rewinding and replaying evolution in controlled environments (Blount, Borland, and Lenski 2008; Fortuna et al. 2013; Losos 1994; Travisano et al. 1995; Desjardins this volume). A recent advance in experimental biology also offers an approach that permits the tape of life to be rewound for individual genes and proteins. Commonly referred as “ancestral sequence reconstruction”, or paleogenetics, this method integrates molecular phylogeny with experimental biology and *in vitro* resurrection of inferred ancestral proteins (Figure 13.1; see also Dean and Thornton 2007). Here I present a novel approach that merges ancestral sequence reconstruction with experimental evolution. In this method, my colleagues and I have engineered a reconstructed ancestral gene directly into a modern microbial genome with the intention of observing the immediate effects of this ancient protein in a modern bacterial context (Kacar and Gaucher 2012). This approach therefore initially follows a backwards-from-present-day strategy in which we essentially recover a previous point on the tape of life, reconstruct it (rewind), and then observe the interaction of this ancient protein with the present (replay). Before presenting the details of our system, and where we would like to go with the *in vivo* resurrection of ancient genes, I will first detail how ancestral sequence reconstruction provides insights into the past.

## Ancestral Sequence Reconstruction: Using Phylogeny to Unravel Evolutionary History

Almost half a century ago, Pauling and Zuckerkandl recognized the potential of phylogenetics to provide information about the molecular history of life. In a pioneering paper titled “Chemical Paleogenetics: ‘Molecular Restoration’ Studies of Extinct Forms of Life,” they suggested a relatively straightforward methodology in which gene or protein sequences obtained from existing organisms would be determined, aligned, and then used to construct phylogenetic trees from which the ancestral states of modern genes and protein sequences could be inferred (Pauling and Zuckerkandl 1963). In 1990, Benner et al. realized the vision of Pauling and Zuckerkandl by reconstructing, synthesizing, and characterizing an ancestral ribonuclease from an extinct bovid ruminant in the laboratory (Stackhouse et al. 1990).

Ancestral sequence reconstruction has since been used to examine the molecular history of many proteins, allowing biologists to test various hypotheses from molecular evolutionary theory by tracing the evolutionary paths proteins have taken to acquire their particular function (Benner, Sassi, and Gaucher 2007). This backward-from-today strategy has also provided helpful insights about various the environmental conditions of the ancient Earth (Galtier, Tourasse, and Gouy 1999; Gaucher 2007; Ogawa and Shirai 2013), paved the way for emerging paradigms such as functional synthesis (Dean and Thornton 2007), evolutionary synthetic biology (Cole and Gaucher 2011), and evolutionary biochemistry (Harms and Thornton 2013). Moreover, it has led to the development of models of molecular evolution that have been used to guide applications of evolutionary theory in medical and industrial settings (Kratzer et al. 2014; Risso et al. 2013).

Our work has focused on reconstructing the molecular past of Elongation Factor-Tu (EF-Tu) proteins. EF-Tu proteins function to deliver aminoacylated-tRNA molecules into the A-site of the ribosome, and are thus essential components of the cell (Czworkowski and Moore 1996). Moreover, EF-Tu proteins also interact with a variety of other proteins functioning outside of translation machinery, and thus serve in activities outside of their role in the ribosome (Defeu Soufo et al. 2010; Kacar and Gaucher 2013; Pieper et al. 2011).

Various ancestral sequences of bacterial EF-Tu, ranging from approximately 500 million to 3.6 billion years old, have been previously inferred and reconstructed (Gaucher, Govindarajan, and Ganesh 2008). Useful to these studies was the information that EF-Tu proteins exhibit a high correlation with it’s organism’s environmental temperature (Gromiha, Oobatake, and Sarai 1999). For example, EF-Tus from thermophilic organisms (organisms living in high temperature environments) exhibit high temperature tolerance and thermostability, while EF-Tus obtained from organisms that live in moderate conditions (i.e. mesophiles) exhibit much lower temperature tolerance and stability. EF-Tus therefore appear to be subject to strong selection to be optimized to their host’s thermal environment.

The combination of EF-Tu’s essential role in the cell, the strong selection acting on EF-Tu phenotype and the availability of various phenotypically and genotypically altered ancient EF-Tu proteins created an ideal situation for us rewind the molecular tape of life for one gene. A rather obvious question is whether a living organism could function if its native EF-Tu was replaced with an ancient form? How and to what degree the reconstructed ancestral EF-Tu proteins would interact with the components of the modern cellular machinery? Moreover, can we provide insights into the factors that determine whether a reconstructed ancient EF-Tu, with very distinct genotypic and phenotypic properties, can interact with modern components?

We therefore set out to first identify whether a recombinant bacteria, in which its endogenous EF-Tu is replaced with an ancestral EF-Tu at the precise genomic location would result in a viable bacteria. First to assess was whether ancient EF-Tu proteins would interact with the members of the translational machinery and thus perform their primary function. For this, we measured the activity of various ancient EF-Tu proteins in an *in vitro* translation system. This system is composed of translation machinery components recombinantly purified from modern *E. coli* bacteria, and individually presented in a test tube, providing us control over which molecules are presented or omitted (Shimizu et al. 2001). We removed the modern *E. coli* EF-Tu from this cell-free system, individually inserted ancient EF-Tu proteins one by one, and measured the activity of the hybrid system by following fluorescent protein production by the translation machinery.

Our *in vitro* findings show that foreign EF-Tus (both ancient counterparts and modern homologs of *E. coli* EF-Tu) exhibit function, albeit not equal to the *E. coli* EF-Tu, in a modern translation system where all the other components of the translation system is obtained from *E. coli* (Zhou et al. 2012). Therefore while replacing the native, modern EF-Tu with an ancient EF-Tu seems likely to put stress on *E. coli,* the recombinant organism is also likely to be viable. Although this is an exciting observation, replacing the EF-Tu of a natural system with another EF-Tu protein that has co-adapted to a foreign cellular system (be it a modern homolog or ancient) will always carry the risk that the interactions between the foreign protein and its ancillary cellular partners may be damaged to the point of complete dysfunction (Figure 13.1). Such a possibility restricts our ability to move genes between organisms; we are fundamentally limited by the historical adaptive mutations and the epistatic relationships that define any particular organism. Before I discuss our success at creating such a recombinant, I will address a question that has no doubt occurred to the reader: Why even bother performing such an experiment with a reconstructed ancestral EF-Tu variant instead of a modern homolog from a different living organism?

**[Figure 13.1].**
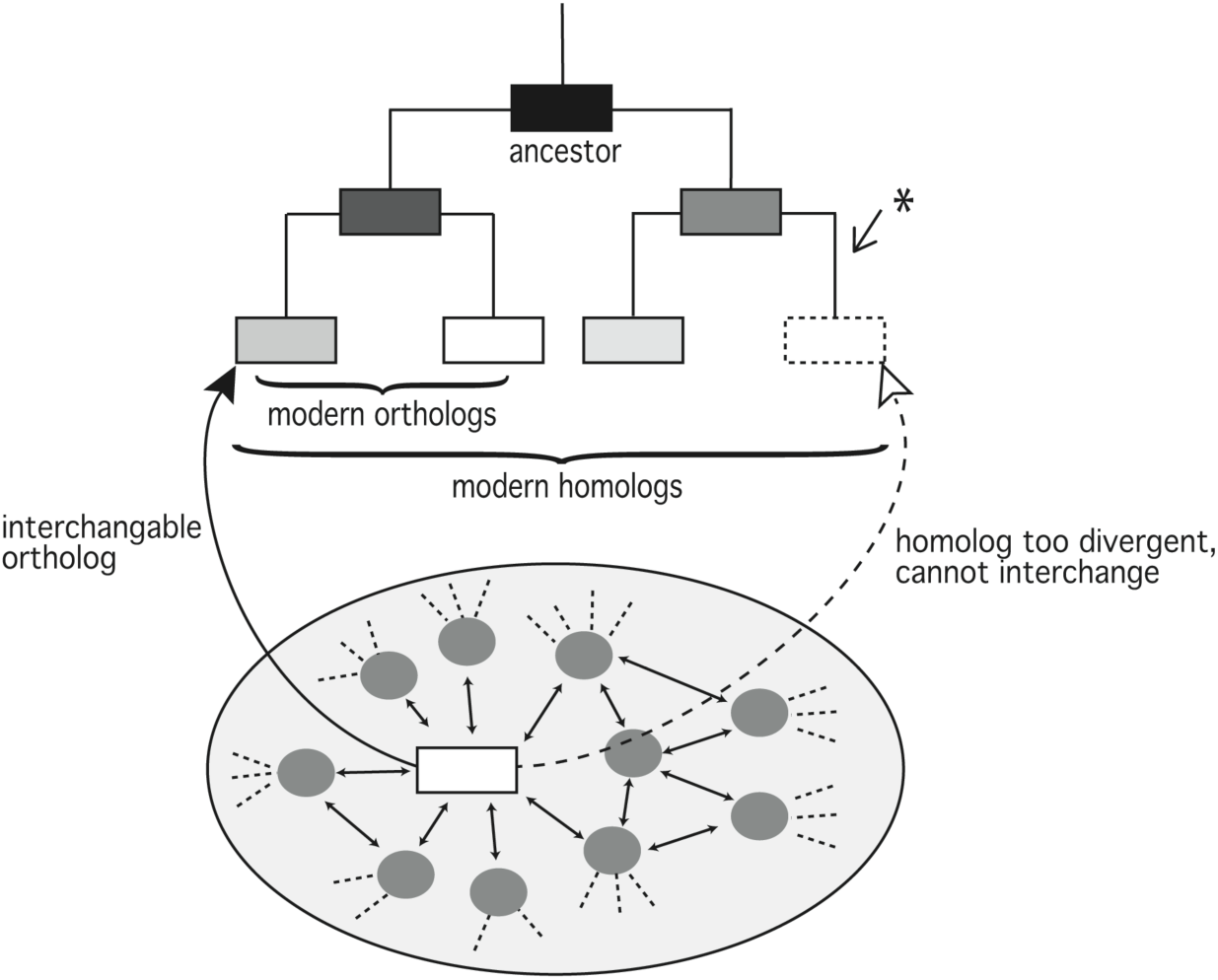

A simplified phylogenetic tree is shown. Phylogenetic trees help us to visualize the evolutionary relationships among organisms. Modern gene and/or protein components are at the tip of the tree. Genes and/or proteins that are connected by a branching point are more closely related to each other and may exhibit more closely related functional properties. In ancestral sequence reconstruction, homologous characters of modern gene or protein components are used to infer the character state of the common ancestor (located at the root of the tree, in black). A central component of a cell (represented with the white box) will exhibit various interactions with other ancillary cellular partners (represented with gray circles) that also have exhibit interactions with many other cellular components (represented with dashed lines). To understand the evolutionary mechanisms that underlie protein function, we need to acknowledge that intrinsic and extrinsic properties of the protein within its interaction network, not just the properties of an individual protein, contribute to and perhaps define how a protein performs its function in a given environment. Replacing a modern gene with its modern homologous or orthologous counterpart from another organism would allow us to identify the functional constraints that shape the two proteins. However, such an approach cannot directly speak to how the replaced protein evolved because its replacement is not on the same line of descent. By replacing a fine-tuned member of a networked system with another component that has co-adapted to another networked system (shown in dashed box), the interactions between the replaced protein and other native proteins may be damaged to the point of producing a non-functional system due to evolved incompatibilities in the homolog (represented by an asterisk, *).]

## Why Engineer a Modern Bacterial Genome with a Reconstructed Ancient Gene?

One common way to assess how a protein may function in a foreign host is do so directly by removing that protein from its host and inserting it into that foreign organism. One might ask why our new approach does not involve replacing a modern gene with a homolog or ortholog from another modern organism. Ultimately, swapping a protein with its homolog has been and will continue to provide valuable insights into functional constraints that shape the two proteins, regardless of whether or not they share functional identity (Applebee et al. 2011; Couñago, Chen, and Shamoo 2006). However, this approach has the drawback that swapped proteins may not share a direct line of descent that connects them over evolutionary time (Figure 13.1). This lack of parity presents the possibility of “functional non-equivalence”, meaning that the two proteins may have traversed two separate and possibly functionally divergent adaptive paths that now prevent the two homologs from functioning properly after being swapped between organisms (Figure 13.1). Moreover, it is important to note that not all of the mutations of the modern homologs are adaptive; random mutations that are results of stochastic events, for instance, can lead to the accrual of mutations in a modern homolog that could prevent the functionality of the modern homolog in another organism once swapped, even while having been neutral under the conditions in which they were accumulated (Camps et al. 2007; Romero and Arnold 2009).

Despite the problems involved, studies in which modern genes are replaced by resurrected ancestral genes are worth the effort. In their article on the mechanistic approaches to study molecular evolution, Dean and Thornton remark that “[t]he functional synthesis should move beyond studies of single genes to analyze the evolution of pathways and networks that are made up of multiple genes. By studying the mechanistic history of the members of an interacting gene set, it should be possible to reconstruct how metabolic and regulatory gene networks emerged and functionally diversified over time” (Dean and Thornton 2007). It is not yet feasible to rewire a whole network but replacing a key component of a complex network with its ancestral counterpart complements these efforts. It is expected that the ability of the interaction network to function with the ancestral component will be limited by incompatibilities that have accrued by the rest of the network over time since the point from which the ancestor was resurrected. A remaining question is how the modern components of a cellular system would co-function with their ancestral interaction partners, if given a second chance.

## An Ancient-Modern Hybrid Test Organism

Following up on our *in vitro* work, we were set to generate a strain of modern *E. coli* harboring an approximately 500 million-year-old EF-Tu protein at the precise chromosomal location of the modern EF-Tu. The ancestral protein is inferred to have been functioning within the common ancestor of γ-proteobacteria and has 21 out of 394 amino acid differences with *E. coli* EF-Tu, and its melting temperature is comparable to *E. coli* EF-Tu (39.5 °C versus 37 °C, respectively).

Marking the first genomic resurrection of a reconstructed ancestral essential gene in place of its modern counterpart within an extant organism, we were able to obtain a viable *E. coli* strain that contains an ancient EF-Tu as the sole genomic copy (Kacar and Gaucher 2012). We next measured the doubling time of the recombinant organism hosting the ancestral EF-Tu. Consistent with our expectations, the hybrid organism exhibits lower fitness, as demonstrated by a doubling time twice as long as its wild type parent strain (Figure 13.2). The hybrid is viable, showing that the ancient EF-Tu can complement essential functions of a descendant in a modern organism. Moreover, the mutations that have accumulated in the components of the modern bacteria over time do not prohibit the modern components that are essential for viability from interacting with the ancient EF-Tu protein. This is an intriguing result, especially considering the vast number of proteins EF-Tu interacts with beyond its primary interaction partners in the translation machinery. The hybrid is also less fit, demonstrating that the ancestral component triggered a stress on the modern organism, creating an ideal case to monitor the co-evolution of the ancient component and the recombinant bacteria.

**[Figure 13.2].**
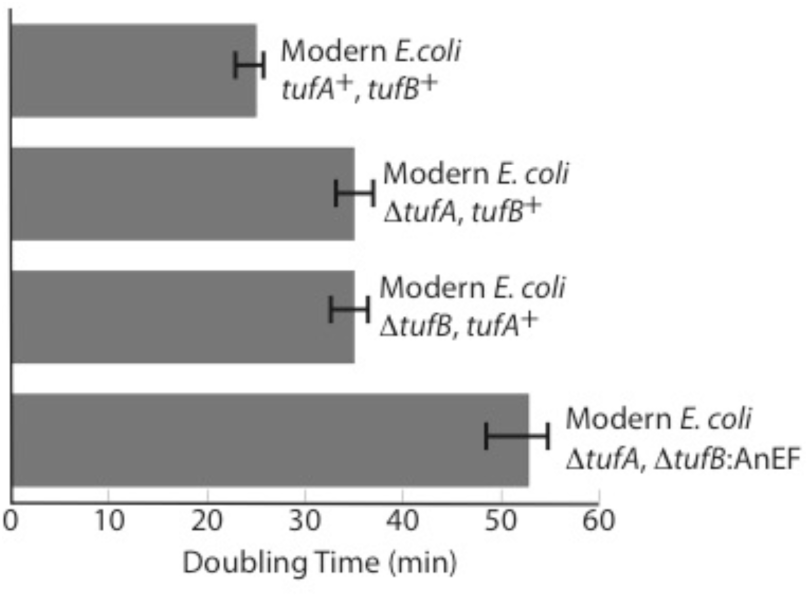

Replacement of a modern essential gene with its ancestral counterpart reduces host bacterium fitness. Precise replacement of a modern bacterial EF-Tu gene with its ~500 million year old ancestor extends the bacterial strain REL606’s doubling time by two-fold. Two genes, *tufA* and *tufB,* (varying by just one amino acid) code for EF-Tu proteins in modern *E. coli.* We deleted the *tufA* gene from the chromosome and replaced the *tufB* copy with the Ancient EF (referred in the figure as AtufA, AtufB:AnEF). Deletions of *tufA* or *tufB* in the *E. coli* REL606 strain have similar effects (~ 34 minutes) when deleted individually. Measurements are performed in rich growth media at 37°C in triplicates. Figure adapted from (Kacar and Gaucher 2012).]

One interesting aspect of this modern/ancient recombinant approach is that it presents a new means of understanding how protein function evolves as a consequence of changes in protein sequence, which in turn may allow us to elucidate how changes in these functions correlate to the overall cellular context within living organisms. Although this is not the primary motivation for setting up these experiments, the approach uniquely complements other studies focused on understanding *in vivo* evolution of proteins. Here I would like to briefly discuss why this peripheral aspect carries biological significance.

Modern cells are the products of immensely long evolutionary histories, and are thus the heirs of a biological heritage that was shaped by the genetic and developmental characteristics of their ancestors, which were themselves shaped by the environments in which they lived. Within this heritage are highly conserved proteins, the functions of which are so crucial that the cellular machinery will not tolerate much change, and which therefore serve as functional fossils. Other proteins are either highly resilient, of less crucial function, or else encode some other flexible potential, and so can readily change in response to exigencies. These proteins show higher rates of evolution, and by definition, lower levels of conservation over time and across taxa. The two groups may be readily distinguished by comparison of proteins across taxa (Baker and Šali 2001; Kominek et al. 2013; Martí-Renom et al. 2000; Papp, Notebaart, and Pál 2011; Wellner, Raitses Gurevich, and Tawfik 2013).

In order to assess how a protein performs its function, a basic approach is to couple molecular biology and biochemistry by removing the protein from the cellular context, and measuring the protein’s activity through its interactions with a defined reactant in a test tube. To experimentally identify what role the *a priori* predicted functionally important sites play, the predicted sites are altered via tools such as site-directed mutagenesis and the mutant protein’s function is measured *in vitro*. Two central points come from this; first, properly predicting the functionally important sites, second, properly analyzing the protein’s function so that studies reflect the protein’s *in vivo,* and thus biologically-relevant, function in the cell.

To address the first point, in order to predict what sites or domains of a protein carry functional importance, one common way is to analyze the evolutionary conservancy of the protein sequence. This method briefly follows the alignment of the amino acid sequences of multiple protein homologs, and then accessing the conserved, different and similar sites through comparison of the aligned taxa. As expected, functionally important sites of a protein will be the most highly conserved across homologs due to their crucial role in protein function (Benner 1989). However, this straightforward methodology does not fully confront the fact that a protein site that is not necessarily conserved across homologous organisms can also carry a functionally crucial role that goes beyond the immediate primary sequence of the protein itself, and may represent an adaptation to a host’s specific intracellular and extracellular environments (Fraser 2005; R. A. Jensen 1976; Khersonsky, Roodveldt, and Tawfik 2006). Recent approaches that include trajectory-scanning mutagenesis and identify regions in which co-evolving proteins interact with each other provide valuable measures addressing this challenge (Ashenberg and Laub 2013; Capra et al. 2010). Indeed, a single amino acid substitution far away from the primary functional site can greatly impact that site’s function (Copley 2003), indicative of various other factors shaping the evolution of protein function.

This observations lead to the second point that needs to be carefully addressed when studying how proteins evolve. In order to reveal how a protein performs its function by biologically realistic means, protein genotype and organismal phenotype need to be directly connected. Engineering strains lacking a particular protein of interest and then reintroducing a homolog or synthetically reconstructed variants of the protein in these mutant strains holds considerable value. Going one step further and observing the co-adaptation between reconstructed proteins and microbial organisms through laboratory evolution would allow us to examine the protein’s function *in vivo.* Finally, this approach would allow us to identify functionally important sites that are not necessarily conserved across taxa but are specific to a host’s intercellular environment, and thus allowing us to consider the context-dependent protein adaptation within a specific lineage.

## Experimental Evolution of Bacteria Containing an Ancient Gene Component

The above discussion provides a context for understanding how functionality is connected to protein divergence. Only after site-specific evolutionary constraints are defined, can we only truly begin to understand how a protein ‘functions’. One alternative way to determine protein functionality is to let an organism mutate its proteome in response to some intra- or intercellular environmental pressure. Such pressures can arise from the availability of novel energy sources or even from the manipulation of internal cellular components. Towards this end, experimental evolution with microbes would provide information on the organismal level by monitoring the real-time evolution of microbial populations to some unique pressure. In these evolution experiments, microbial populations are evolved in the lab through either continuous culture or serial transfer under controlled conditions, permitting evolution to be studied in unprecedented detail in near real time (Elena and Lenski 2003).

The experimental evolution approach is particularly powerful because of the high level of control it permits. Samples of evolving populations may be frozen at regular intervals while still remaining viable, providing “frozen fossil records” that be analyzed to examine a variety of important questions (Elena and Lenski 2003). Experimental evolution may be used for highly detailed study of a variety of questions, such as whether evolution follows contingent or deterministic paths, whether mutations accumulate neutrally or adaptively, and how epistatic interactions impact evolution. Indeed, laboratory evolution experiments using microbes have provided deep insights into the interplay between contingency and deterministic processes in evolution (Blount, Borland, and Lenski 2008; Blount this volume; Losos 2011; Travisano et al. 1995; Vermeij 2006).

The experimental evolution approach is key in our next step with the ancient-modern hybrid organism we have constructed to replay the tape. We are currently evolving the reconstructed organism in the laboratory to study how compensatory evolution alters the ancient gene and members of its epistatic network. Of particular interest is how much and how quickly the construct’s fitness and phenotype will change, and how much of these changes are mediated by the ancient EF-Tu’s direct accumulation of beneficial mutations versus their accumulation in other, associated genes. Moreover, we wish to assess the extent to which the mutational trajectory is guided by random and unpredictable events (i.e., attributable to chance) and how much it is determined directly by the genotype and the phenotype of the ancient EF-Tu engineered inside the bacteria (i.e., attributable to contingency).

Evolving a bacterium containing a single ancient gene in its genome is analogous to replaying a particular track on the tape of life within the context of the modern organism. To play with the metaphor a bit, we made a mix-tape by splicing a very old track from the tape of life, and are now playing it on a modern tape deck (the modern organism). Both the tape and the tape deck can evolve in this case, and we will be able to examine how they evolve and adapt to each other, potentially recapitulating some of the evolution that occurred in the first place. While this sort of replay experiment is not exactly what Gould had in mind, how the organism adapts to the ancestral protein and vice versa promises to shed light on the molecular evolution that took place during EF-Tu evolution.

This approach does have limitations as well. It is important to recognize that the ancient component resurrected in a modern organism’s adaptation will be shaped by its interactions with modern components adapted to both its modern counterpart, and a cellular environment that is millions of years from ahead of the conditions of the ancient component. It is also important to note that the environmental conditions in the laboratory are unlikely to be comparable to the conditions and circumstances under which the ancestral protein evolved into its modern descendant.

Despite its limitations, re-evolving ancient genes in modern organisms permits us to investigate historical aspects of repeatability and parallelism in evolution that replacing a gene with a modern homolog does not. Multiple aspects of the evolutionary processes, including mutation and genetic drift, are inherently stochastic, which makes it challenging to predict how a system hosting a reconstructed gene (as well as a modern homolog) would co-evolve. By setting up laboratory evolution experiments of microbial organisms that carry an ancient component in replicate populations, we are able to observe whether or not multiple populations carrying the ancient component evolve the same way, or in parallel ways, under identical environmental conditions. Engineering constructs with genes encoding different reconstructed ancestral states of the same protein (e.g., 500 million-year-old, 1 billion-year-old, etc.) would permit investigation of whether or not evolutionary trajectories are strictly dependent on evolutionary starting points (Figure 13.3; see also Travisano et al. 1995). For instance, certain adaptive zones may readily be more accessible as we go further back in time; i.e., the older the component we resurrect, the greater the likelihood of finding a mutational point that is capable of functional innovation by resetting the epistatic ratchet (Bridgham, Ortlund, and Thornton 2009).

**[Figure 13.3]:**
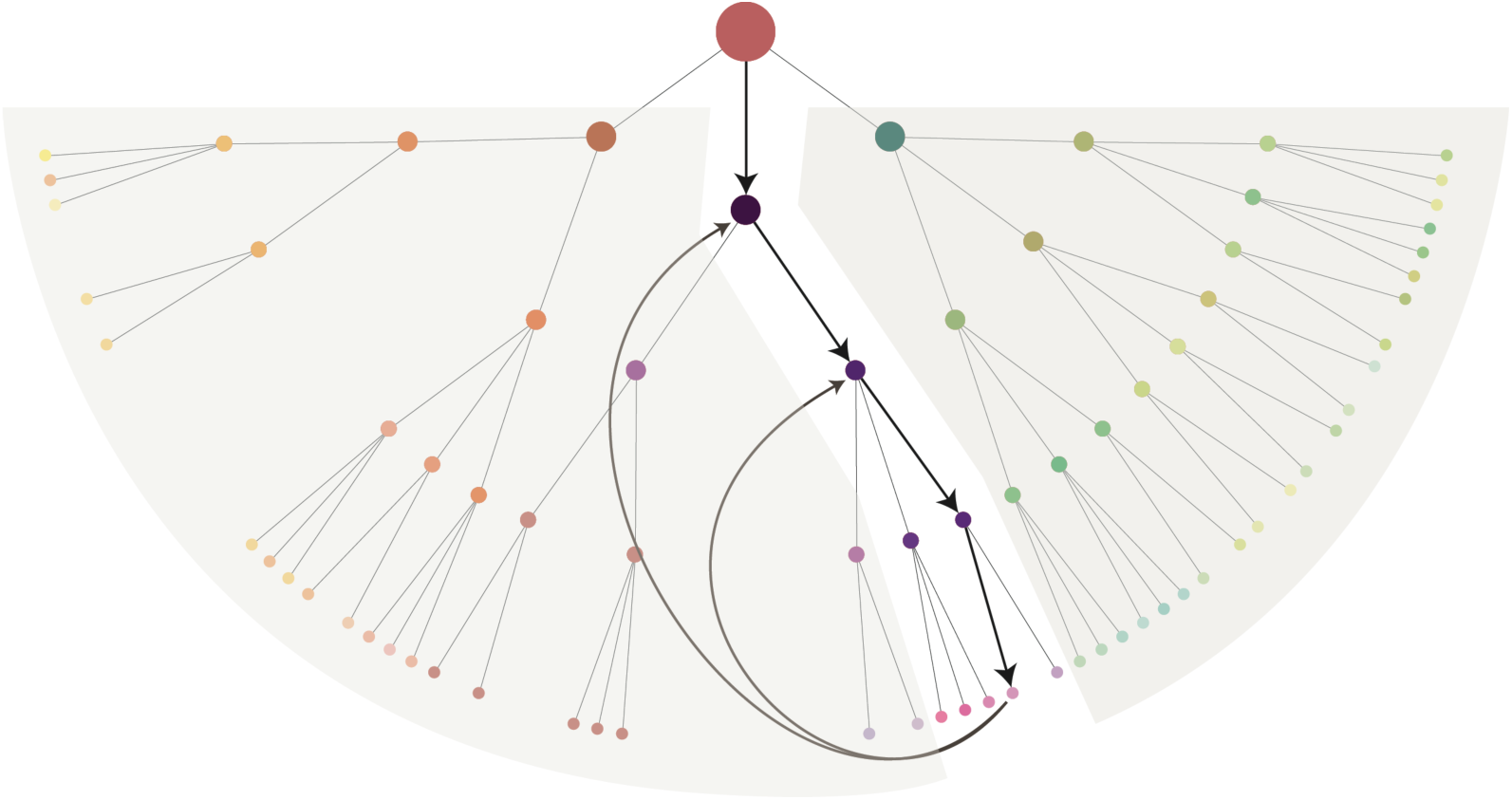

A hypothetical scheme of evolutionary patterns: Interaction between an organism containing ancestral variants of a protein (represented by different colored circles) and its environment will produce a series of adaptive zones. Replacing a modern protein with an ancestral counterpart may potentially lead the organism to explore a wider potential adaptive landscape. Replaying evolution starting from successively older time-points (shown with successively longer arrows) would then allow access to points corresponding in time to historic evolutionary states that preceded the accumulation of mutational steps, thus allowing the engineered organism to operate in different adaptive zones.]

## Concluding Remarks

To directly examine the relationship between historical constraints and evolutionary trajectories, and to assess the role historical contingency played in shaping protein evolution within a biologically realistic framework, it is important to determine the historical paths of proteins, both genotypic and phenotypic levels, and then study how these paths affected the organism. Many properties of protein-protein interaction networks, not just those of individual proteins, contribute to and perhaps define how a protein performs its function in a given environment. Consequently, the biochemistry of proteins may be intimately linked to their host organisms’ behavior.

The only lesson history teaches is that no one learns from history (Hegel 1953). This playful quote is not the case for understanding the role history plays in shaping evolution. Integrating molecular-level theory into studies of evolutionary history and merging ancestral sequence reconstruction studies with experimental evolution will help to provide mechanistic explanations into several long-standing questions in evolutionary biology: How does an organism’s history shape its future trajectories, and are there deterministic paths along these trajectories? Does evolution inevitably lead to the same endpoints? How much does the past restrict the future? Moreover, how do answers to these questions depend on the genetic and the environmental conditions of the system examined? Synthesis of the fields of synthetic biology, biochemistry, and experimental evolution holds great promise for generating a new understanding of functional, structural and historical constraints that shape biological evolution. Indeed, this synthesis holds great promise for learning much of and from life’s history.

## Acknowledgements

This work was supported partly by the NASA Astrobiology Institute through a NASA Postdoctoral Program Fellowship, a NASA Exobiology and Evolutionary Biology grant NNX13AI08G and a NASA Astrobiology Institute Early Career Research Award. Eric Gaucher, Zachary Blount, W. Seth Childers and Zach Adam provided helpful comments and suggestions on an earlier version of this chapter, Georgia Tech undergraduate students Lily Tran and Jen Zhang provided invaluable assistance in the laboratory, Rich Lenski and Neerja Heala (MSU) provided the *E. coli* REL606 and REL607 strains.

